# Optimized structure of monoubiquitinated FANCD2 (human) at Lys 561: a theoretical approach

**DOI:** 10.1101/2021.03.12.435201

**Authors:** Sudipa Mondal, Subba Reddy, Sudit S. Mukhopadhyay

## Abstract

Fanconi anaemia pathway repairs inter-strand cross linking damage (ICL) of the DNA. Monoubiquitination of FANCD2 and FANCI is very crucial for ICL repairing. In this work we have tried to understand the monoubiquitinated FANCD2 structure, which facilitates the FANCD2 for binding the damage part of the chromatin. Crystal structure of the monoubiquitinated FANCD2 alone is not available, therefore we have developed the optimized structure of the human monoubiquitinated (Lys 561) FANCD2. As there is no suitable software or web server we have developed a method for building up monoubiquitinated product and validated on simplest monoubiquitinated protein, diubiquitin. We have predicted the structure of human monoubiquitinated FANCD2 by using our method and studied the interaction with DNA by docking studies. Molecular Dynamics (MD) simulation was used to understand the stability of the structure. Large structural differences have been observed between FANCD2 and monoubiquitinated FANCD2. DNA docking studies suggest that the binding site varies for the FANCD2 and monoubiquitinated FANCD2.

## 1. Introduction

Fanconi anemia pathway (FA pathway) is an essential tumor-suppressive pathway (Ceccaldi, et al., 2016; Rodríguez et al., 2017) and it is required for protecting the human genome from inter crosslink (ICL) DNA damage. Till today, twenty two (22) genes (FANCA, FANCB, FANCC, FANCD1/BRCA2, FANCD2, FANCE, FANCF, FANCG, FANCI, FANCJ/BRIP1, FANCL, FANCM, FANCN/PALB2, and FANCO/RAD51C, FANCP/SLX4, FANCQ/ERCC4, FANCR/RAD51, FANCS/BRACA1, FANCT/UBE2T, FANCU/ XRCC2 FANCV, FANCW) have been identified which are involved in the FA pathway (Bhattacharjee & Nandi, 2017; Deakynel & Mazin, 2011; Joshi et.al, 2019). Eight of the FA proteins (FANCA, FANCB, FANCC, FANCE, FANCF, FANCG, FANCL, and FANCM) form a nuclear core complex (FA complex), which activates the FA pathway through regulation of the monoubiquitination of FANCD2 and FANCI (Walden & Deans, 2014).

The most crucial event in the FA repair pathway is the monoubiqutination of FANCD2- FANCI complex (Zang et al., 2007; Ulrich & Walden, 2010). Upon monoubiqutination the FANCD2 recognises the damage part of the chromatin and interact with damage part for repairing along with other repairing protein complex. Monoubiqutination of protein is a posttranslation modification which regulates many cellular processes (Glickman & Ciechanover, 2002; Mukhopadhyay & Riezman, 2007; Schnell & Hicke, 2003). Many important proteins require monoubiqutination for activation and execution of cellular function (Chen et al., 1996). Many proteins have found to alter their stability after addition of single ubiquitin moiety (Hagai & Levy, 2010). In the FA pathway the FANCD2 requires monoubiqutination for recognition and binding of the damage part of the DNA, but how a single ubiquitin influence the FANCD2 protein or gives the stability is not very clear. As there is no software available to carry out the monoubiquitination theoretically, we have developed a method for calculating the arrangement of monoubiquitinated product and then have predicted the monoubiquitinated FANCD2 structure by using our methodology. The result have been hypothesized that the single ubiquitin moiety alters the physical structure which facilitates the FANCD2 for recognition and binding to the specific damage part of the chromatin. We have also found that the DNA binds to FANCD2 and monoubiquitinated FANCD2 at different sites.

## 2. Experimental

### Methods

#### 2.1. Homology modelling

Crystal structure of human FANCD2 was not available in the Protein Data Bank so homology modelling using MODELLER (Webb & Sali, 2016; Marti-Renom et al., 2000; Sali & Blundell, 1993; Fiser et al., 2000) was done to predict the 3D structure of human FANCD2. Human FANCD2 is a 1,451–amino acid (164-kDa) protein, the sequence of the FANCD2 protein was downloaded from NCBI database in FASTA format. Crystal structure of mouse FANCD2 (PDB is 3S4W) was taken as template which showed 77% sequence identity by BLAST search. Modelled structure was crossed checked by doing further modelling using I-TASSER web server (Roy at al., 2010; Yang et al., 2015; Yang & Zang, 2015). The generated modelled protein was energy minimized by steepest descent method to avoid any stereo chemical restraints. The model was further validated by Ramachandran plot, Verify3D (Eisenberg et al., 1977) and ProQ (Cristobal et al., 2001).

#### 2.2. Protein-protein Docking

Protein-protein docking was done using Dock Protein toolbar under the macromolecule module of the Discovery Studio software (BIOVIA-Discovery Studio, 2017). The ZDOCK protocol was used to carry out rigid body docking of two proteins that were developed using pair wise shape complementarity, desolvation, and electrostatic energies (Chen et al., 2003). The program employs a fast Fourier transform (FFT) correlation technique that is used to explore the rotational and translational space of protein–protein system. A total of 54,000 structures or poses were generated by grid searching. The rotational space was 6°. The top 2000 poses were retained for evaluation using ZRANK score (Pierce & Weng, 2007) and processed by the clustering method. The poses with the lowest ZRANK score was selected for further MD simulations. The final structure was evaluated by Ramachandran plot, ProQ and Verify3D. Solvent-accessible surface area (SASA) study was used to know the buried surface area of the FANCD2 protein.

#### 2.3. Molecular dynamic simulation

Molecular dynamic simulation is important to predict the dynamic property and the optimized structure of a protein (McCammon et al.,1980; Zhang at al., 2009; Freddie & Salsbury, 2010). MD simulations mimic the physical motions of atoms in the protein molecule present in the actual environment. The atoms are allowed to interact for a certain period of time, which will help to compute their trajectory in and around the protein molecule. MD simulation was done using Charmm (Brooks et al., 2009) force field using Run simulation toolbar under the Simulation module of the Discovery Studio software package. Before the simulation run we solvated the protein with water molecule in a explicit periodic boundary (orthorhombic cell shape). The system was dissolved in TIP3P water molecules (Jorgensen et al., 1983). The solvated protein was first minimized using 1000 cycles steepest descent followed by a second minimization using conjugate gradient by 2000 cycles (Chaudhuri et al, 2019). The minimization was done to remove any unfavourable contact here in the structure. Next the minimized structure was heated from 50K to 300K for 120ps followed by the equilibration in NVT [constant number (N), volume (V), temperature (T)] and NPT [constant number (N), pressure(P), temprature (T)] ensemble for 1000ps each. Long range electrostatic interaction were calculated using Ewald particle mesh method (Cheatham et al., 1995). After equilibration the protein structure was simulated for 20ns at 300K with a time step integration for 2fs for dynamic production run. This was carried in the NVT ensemble. During the simulations, all covalent bonds involving hydrogen were constrained using the SHAKE algorithm (Ryckeart et al., 1977).

#### 2.4. DNA- Protein docking studies

Docking is an important tool to study how two molecular structures fit together and to provide mechanistic insight (Mondal et al., 2015). Docking studies of DNA with FANCD2 and monoubiquitinated FANCD2 was done using PatchDock web server (Schneidman-Duhovny et al., 2005). The PatchDock method performs structure prediction of prote-protein, protein-small molecule, DNA-small molecule, DNA-protein complexes (Mondal et. al., 2018; Pagadala et al., 2017). PatchDock is a geometry-based molecular docking algorithm (Duhovny et, al., 2002), it is aimed at finding docking transformations that yield good molecular shape complementarity.

### Result and Discussion

#### 3.1. Structure generation and optimization of structure of human FANCD2

To establish the three dimensional (3D) structure of human FANCD2 homology modelling has been performed taking mouse FANCD2 as template (PDB:3S4W) structure. Mouse FANCD2 has showed 77% of homologous identity with human FANCD2 when the sequence alignment of protein domain has been performed by Protein BLAST search. The 3D structure has been produced by MODELLER implemented in DS 2017. The best model has been selected from resultant several structures on the basis of lowest probable density function and DOPE (discrete optimized potential energy) score. Stereo chemical property of the final structure has been evaluated by Ramachandran plot and it showed that 97.4% of the residues are in allowed region. To cross check the quality of the modelled structure the sequence has been submitted to I-TASSER server to get the predicted three dimensional structure. The structure with highest C score has been selected as best model and 93% of the residues are in allowed region in Ramachandran plot. Both structures have been compared and it has been found that they are very similar **(Fig 1a, 1b).** LG score value of ProQ results of modelled human FANCD2 (modelled by Discovery studio and I-TASSER) are comparable with that of mouse FANCD2 (crystal structure) (Fig. 1c) which suggests that the model quality is in the range of scores typically found for similar size proteins.

**Fig. 1:**
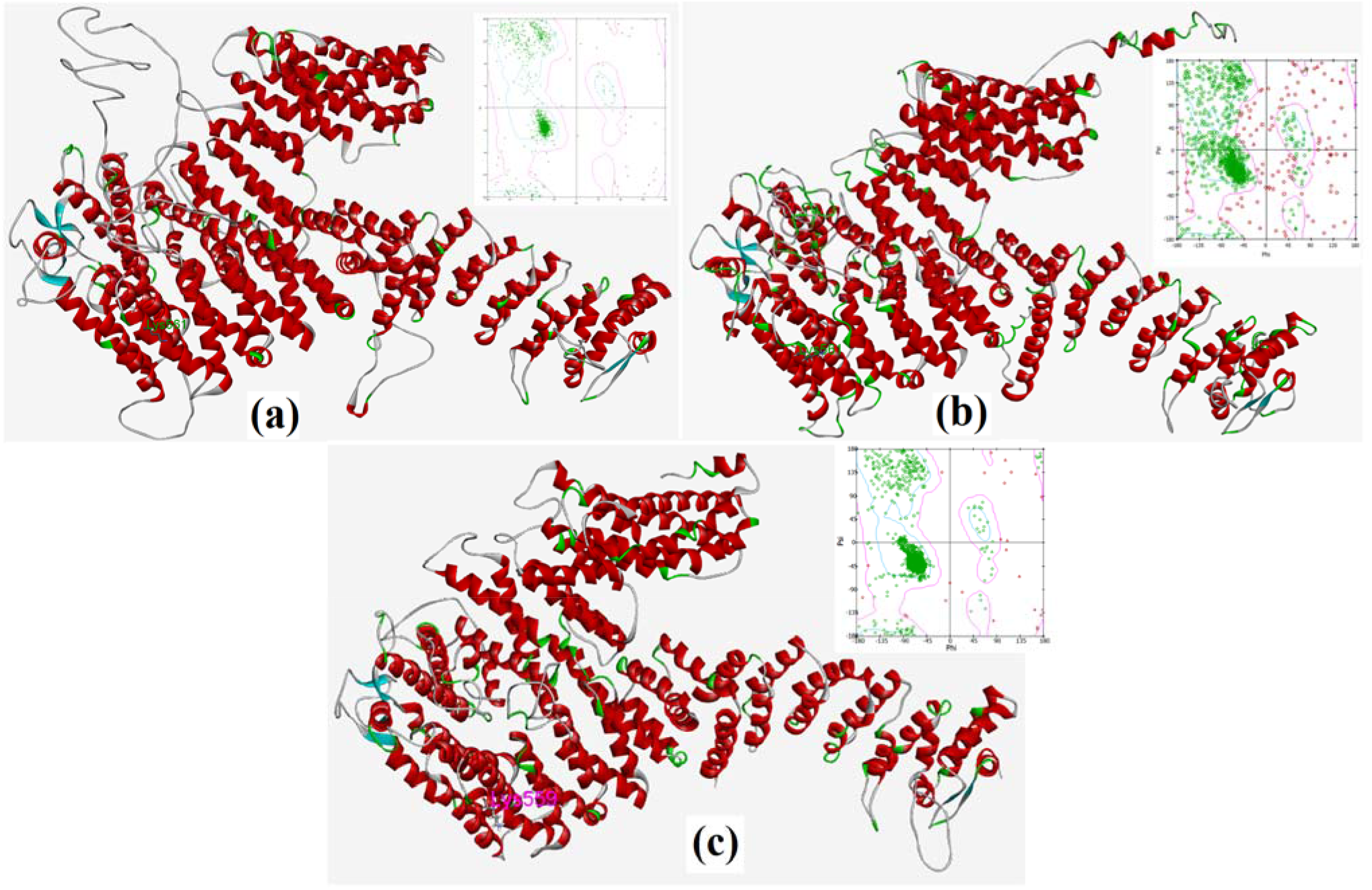
Structure of human FANCD2 modelled by (a) Discovery Studio, (b) I-TASSER server and (c) mouse FANCD2 from crystal structure.

The energy minimized modelled structure has been put for molecular dynamic simulation for 20ns. Potential energy, back bone root mean square deviation (RMSD), backbone root mean square fluctuation (RMSF), Radius of gyration (Rg) have been analysed throughout the trajectory to investigate the dynamic behaviour and stability of the simulation system. After optimization the average value of the potential energy of human FANCD2 is −465500 Kcal/mol. The potential energy profile of the system decreases rapidly in first 1ns and have remained stable throughout the simulation run (**Supplementry file, Fig.S1a**). The backbone RMSD profile of the protein has been compared to their initially minimized structure. The backbone RMSD of the system have shown little deviation in initial conformers **(Supplementary File Fig.S1b)** from 0 to 3 ns and have remained constant thereafter. Root mean square fluctuation (RMSF) has also been analysed (**Supplementry file, Fig. S1c**), no significant fluctuation has been observed among the residues near Lys561 the residue of our major interest, but higher fluctuation is observed for the amino acids near C-terminus and N-terminus. RMSF value during simulation run indicates that both the C-terminus and N-terminus of the FANCD2 are largely flexible. The radius of gyration (Rg) is defined as the distribution of atoms of a protein around its axis. It is the measure of compactness of the protein. The length that represents the distance between the point when it is rotating and the point where the transfer of energy has the maximum effect gives Rg. It is measured as the root mean square distance of the collection of atoms from their common centre of gravity. FANCD2 has maintained a regular value of Rg (60.7± 0.2) Å (**Supplementry file, Fig**. S2) throughout the simulation so we can say that the protein is stably folded. The quality of the final structure **(Fig. 2)** has been evaluated by Ramachandran Plot and it has been observed that 98% of residues remain in the allowed region.

**Fig. 2:**
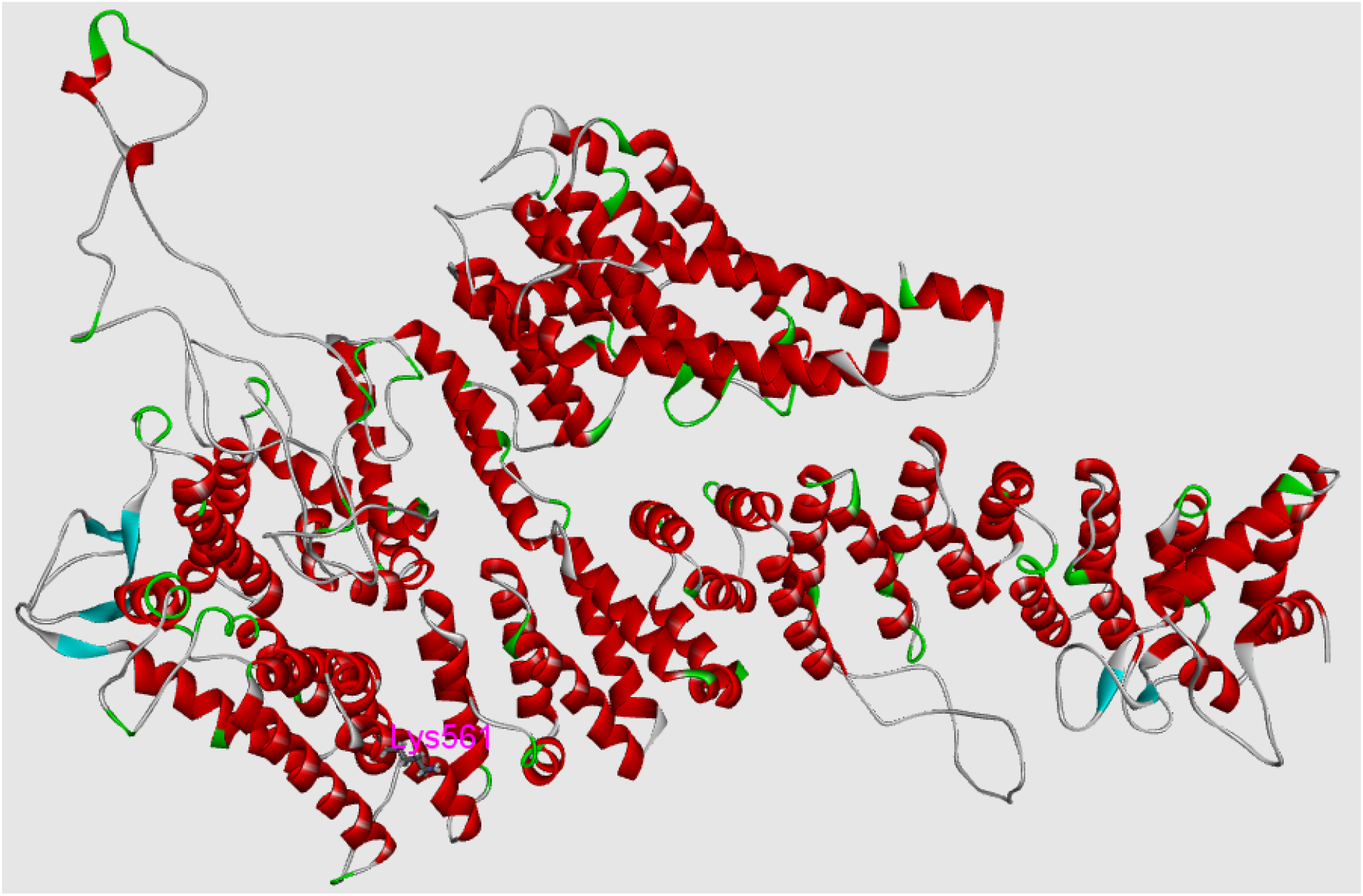
Optimized structure of human FANCD2 after molecular dynamic simulation run.

#### 3.2. Monoubiquitination of human FANCD2

Our aim is to understand the structure of monoubiquitinated FANCD2 protein (human) and to evaluate what structural difference is taking place after the monoubiquitination. Monoubiquitination is a post translational modification where a small protein named ubiquitin is added covalently to the substrate protein. Here the substrate protein is FANCD2 (human) and the monoubiquitination takes place at Lysine 561 position. But major challenge is to establish the structure of monoubiquitinated (at Lys 561) FANCD2 since no software or server do the monoubiquitination directly though there are servers to do other post translational modification like hydroxylation or phosphorylation. So we have made a hypothesis to get the structure of the monoubiquitinated protein and have taken diubiquitins as model system to validate the prediction. The simplest monoubiquitinated proteins are diubiquitins which are basically mono ubquitinated ubiquitins. Ubiquitin have seven lysine residues that serve as point of mono ubiquitination and produces seven dissimilar diubiquitins. Crystal structure is available for all seven diubiquitins (few are of human and few are of cattle). Crystal structure for human system is available for K6, K11and K63 linked diubiquitin (Virdee et al., 2010; Bremm et al., 2010; Komander et al., 2010), whereas the crystal structure available for K29, K33 and K48 linked diubiquitin are of cattle (Kristariyanto et al., 2015 (1); Kristariyanto et al., 2015 (2); Trempe et al.,2010]. NMR structure is available for human 27 linked diubiquitin (Castan eda et al., 2016) [42]. We have predicted the structure of different diubiquitin theoretically and compared that with the reported crystal structure. We have noticed that one thing is common in all the crystal structures; in all diubiquitins terminal glycine residue (Gly76) is attached covalently with lysine residue at different position (Lys6, 11, 27, 29, 33, 48, 63) via an isopeptide bond between –COOH group of Gly76 and ε-(epsilon) amino (-NH_2_) group of respective lysine. To predict the desired structure at first we have done protein-protein docking so that Gly 76 remain in close proximity with the required lysine residue (to do such assortment we have put the filtration parameter while giving input for protein-protein docking) followed by the selection of those poses which has higher Z-score value. Among the selected poses we have made the further refinement by running Rdock (Li et al., 2003] [43] and finally selected that pose which has lowest ER value along with lysine nitrogen (-NH2 of monoubiquitination site) and carboxyl group (-COOH) of Gly 76 are within bonding distance (~1.5Å). It was observed that the selected pose has given rise to maximum stabilization when the amide bond has been made manually using Sketch molecule protocol inside the Small molecule module of Discovery studio between Gly76 residue of one ubiquitin and particular Lys residue of another ubiquitin. Then we have run the molecular dynamic simulation till the energy of the system and RMSD have become constant. The final structure has been compared with the reported crystal structure (PDB id 2XK5, 2XEW, 5UJL, 2JF5, 3M3J) and it has been observed that they are well comparable.

Here it is being described in details how we have predicted the structure of K6-linked diubiquitin and how close it is to the reported crystal structure. We have docked one human ubiquitin protein chain (at Gly76 position) with another human ubiquitin protein chain at Lys 6 position. All poses where Gly 76 and Lys 6 are in close proximity have been put for further refinement by RDock run. Next the poses obtained after RDock have been observed very carefully and selected ten best poses where the carboxyl group (-COO^-^) of Gly76 and amino group (-NH_2_) of the Lys 6 are within bonding distance (~1.5Å) and have high ZDock Score (Supplementary file Table S1). Then we have made the isopeptide bond between the carboxyl group (-COO ^-^) of the one ubiquitin’s glycine (Gly76) and epsilon (ε) amino group (NH_2_) of the lysine at 6 position (Lys 6) of the other ubiquitin. Finally we have chosen that structure which is most stable (minimum energy) after the bond formation. Next the quality of the predicted protein structure (6 linked diubiquitin) has been evaluated by Ramachandran Plot, ProQ, Verify3D (**Supplementary file, Table S2**). For K6-linked diubiquitin it has been observed that 95.97% of the amino acid residues are in allowed region (**Supplementary file Fig. S3**), LGscore in ProQ is 3.77 which indicates it as a very good model structure. After that the molecular dynamic simulation has been run to get the optimized structure. The simulation have been carried on till the energy and the RMSD have reached a constant value. Final arrangement obtained after the MD simulation has been compared with the crystal structure of K6-linked diubiquitin (PDB id 2XK5). It is observed that the theoretically calculated K6-linked diubiquitin is very much similar (**Fig. 3**) with the reported crystal structure since the RMSD after superimposition (Fig.Sxx) is 4.9. Similarly we have predicted the structure of K11, 2K7 and K63-linked diubiquitin of human system and K48-linked diubiquitin of cattle system and have compared with the reported crystal structure (PDB id 2XEW, 5UJL, 2JF5 and 3M3J respectively) (**Fig. 4 and Supplementary file Fig. S4, S5**). All the theoretically calculated molecular arrangements are very near to experimental structure and the corresponding RMSD values obtained after superimposition are 3.2, 4.3, 6.4 and 1.6 respectively. Quality of all the theoretically predicted structures has been verified by Ramachandran plot, ProQ and Verify 3D (**Supplementary file Fig.S6, Table S3**). In Ramachandran plot more than 95% of the amino acid residues are in allowed region. ProQ data also says that all these are good model structure since LG score in all the cases are more than 2.5, in Verify3D tool more than 95% residues lie in the score >2.0 (in Verify3D for good score at least 80% of the residue should lie in the score 2.0). Apparently it seems that secondary structures are missing in the computed diubiquitin proteins as per the pictures in Fig. 3a, 4a, 7a, 8a, 9b, S4a, S5a but if Ramachandran plot (**Fig. 5, S6**) of the computed structures are noticed, it is observed that the residues are lying on proper position related to secondary structures (Wiltgen, Encyclopedia of Bioinformatics and computational Biology, 2019; Laskowski, Swaminathan Comprehensive Medicinal Chemistry II, 2007).

**Fig. 3:**
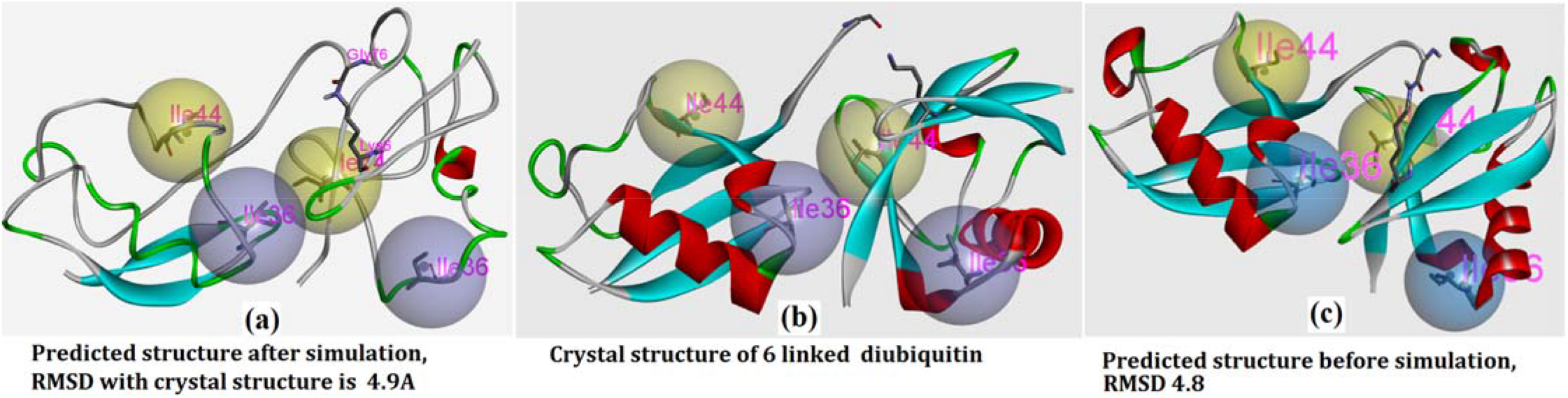
Comparison of predicted 6 linked diubiquitn structure [(a) after MD simulation, (c) before simulation] with the reported crystal structure (b).

**Fig. 4:**
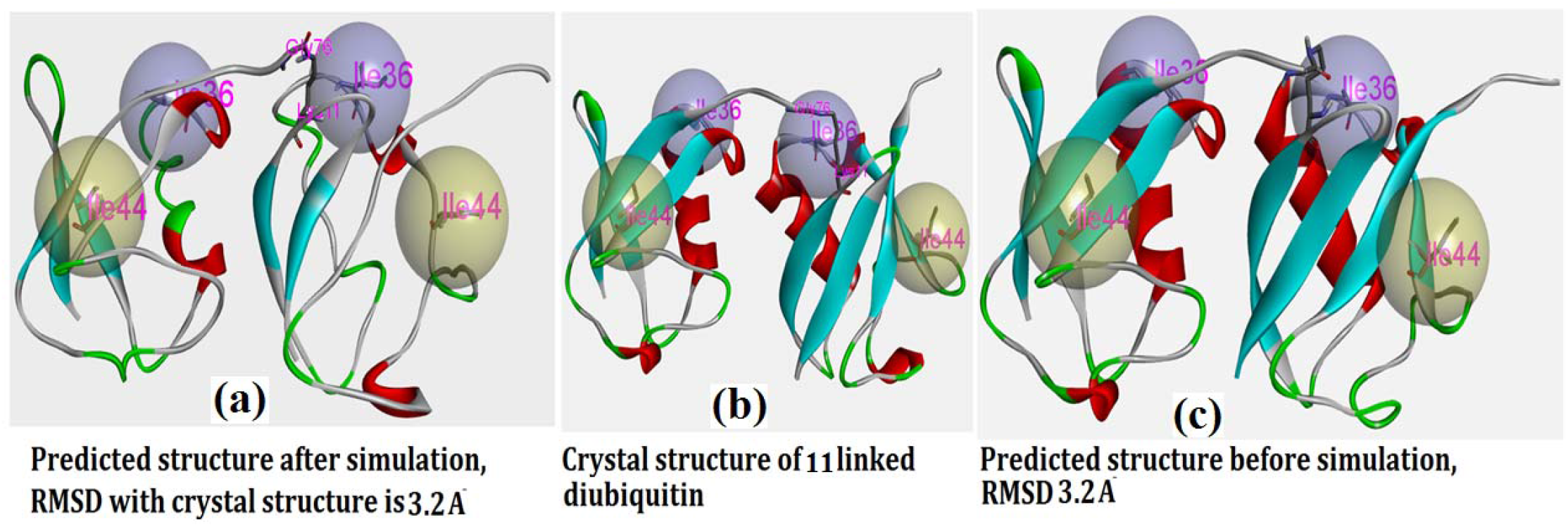
Comparison of theoretically calculated 11 linked diubiquitin (a and c) with its crystal structure (b).

**Figure 5:**
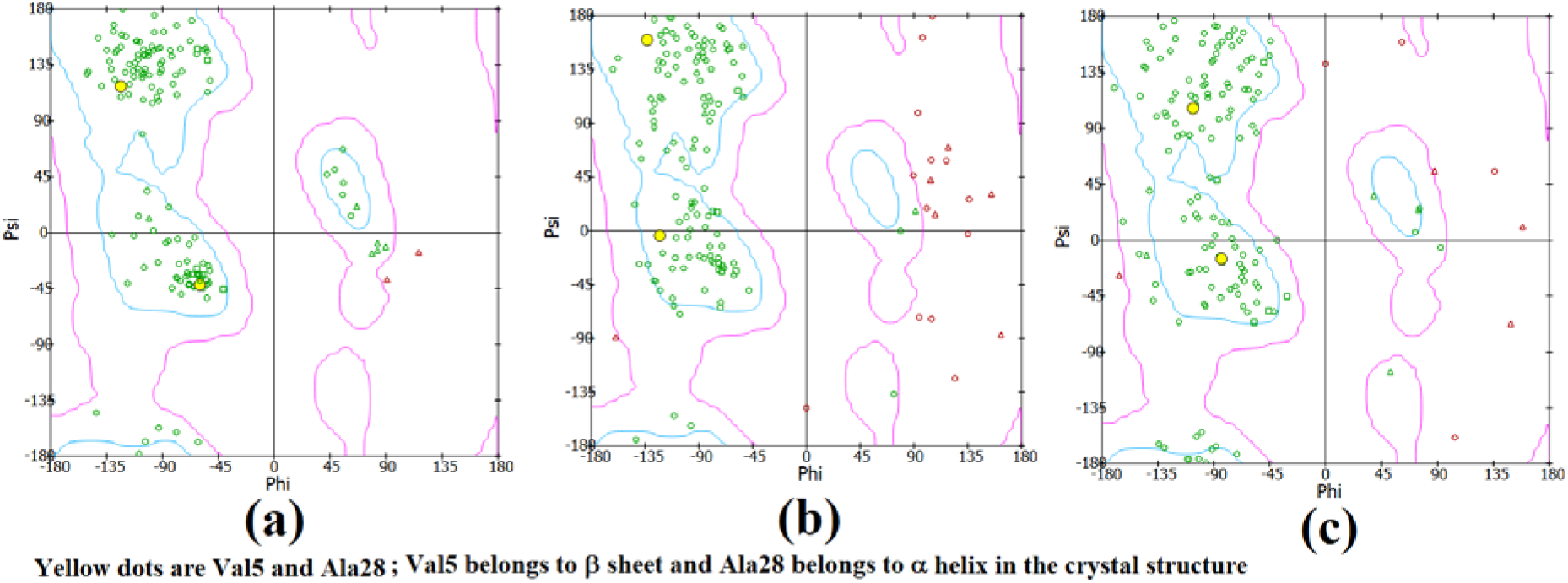
Ramachandran plot of K6 diubiquitin; (a) crystal structure, (b) average structure after simulation, (c) after dynamics equilibration.

We have applied the same steps on human FANCD2, first protein-protein docking has been done at Lys 561 position of FANCD2 with the Gly76 position of human ubiquitin (PDB id 1UBQ) by ZDock, then refinement of the resulted poses by RDock followed by the isoamide bond formation between -COOH of Gly76 and -NH_2_ of Lys 561 in those poses where they are in bonding proximity and finally selection of that pose which has maximum ZDock score **(Table 1**) and highest stabilization have taken place due to bond formation. The resultant structure **(Fig. 6)** has been evaluated by Ramchandran plot, ProQ and Verify3D. 97% of the residue of the monoubiquitinated FANCD2 are in allowed region, the LG scores in ProQ is more than 2.5 which indicates it as a very good structure. Verify3D points it as a very good structure as 97% of the residue lie in the score 2.0.

**Fig. 6:**
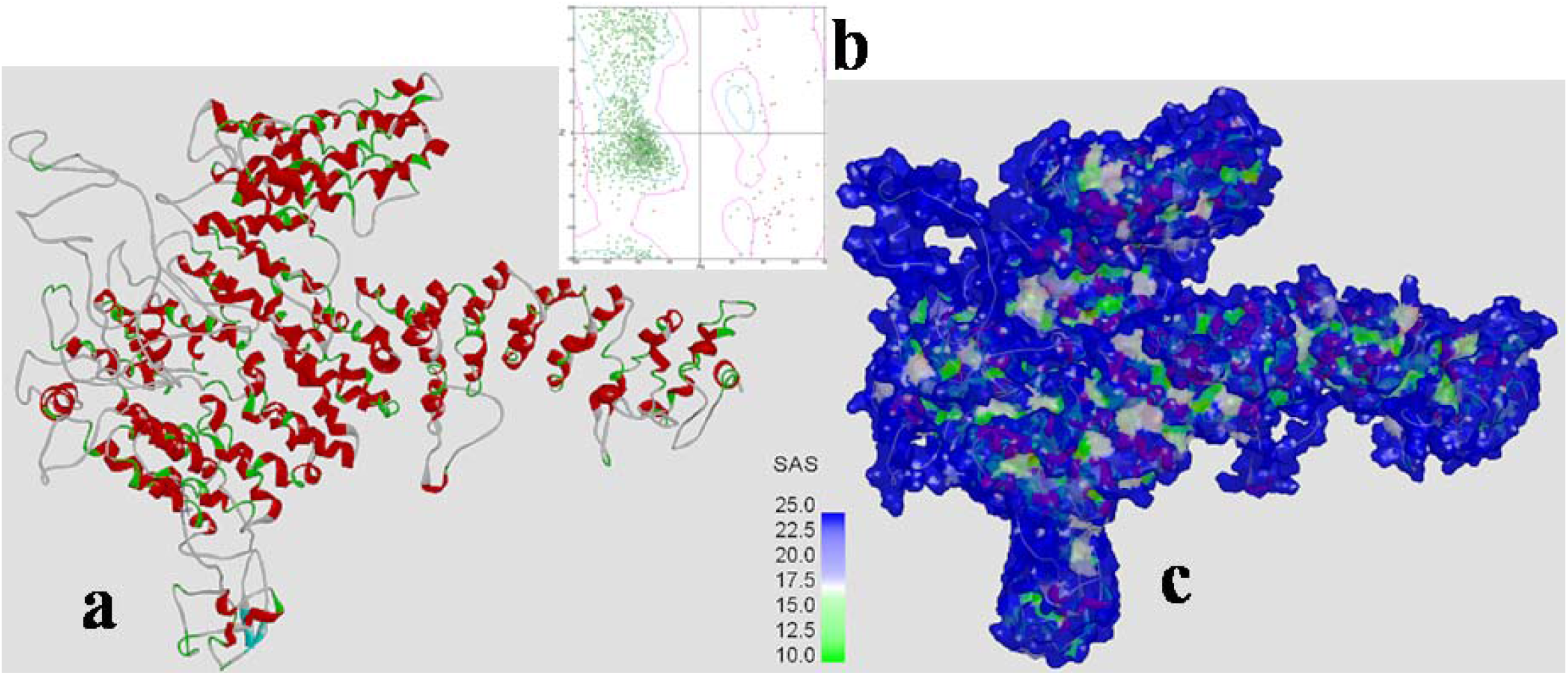
Structure of monoubiquitinated FANCD2 at Lys561 position (a), Ramachandran plot (b), surface picture (c) before simulation.

**Table 1:**
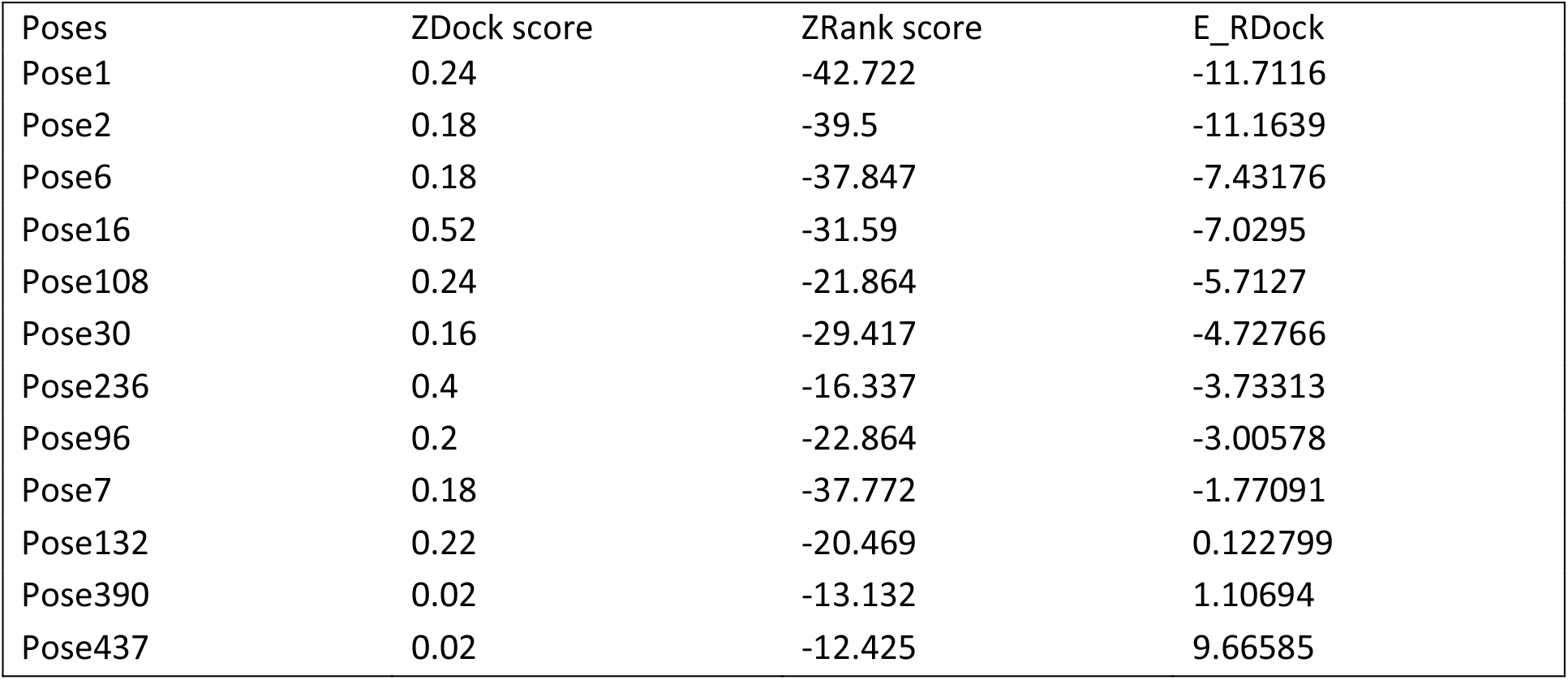
Data obtained when ubiquitin (at Gly76 position) is docked with human FANCD2 at Lys561 position.

| Poses | ZDock score | ZRank score | E_RDock |
| --- | --- | --- | --- |
| Pose1 | 0.24 | -42.722 | -11.7116 |
| Pose2 | 0.18 | -39.5 | -11.1639 |
| Pose6 | 0.18 | -37.847 | -7.43176 |
| Pose16 | 0.52 | -31.59 | -7.0295 |
| Pose108 | 0.24 | -21.864 | -5.7127 |
| Pose30 | 0.16 | -29.417 | -4.72766 |
| Pose236 | 0.4 | -16.337 | -3.73313 |
| Pose96 | 0.2 | -22.864 | -3.00578 |
| Pose7 | 0.18 | -37.772 | -1.77091 |
| Pose132 | 0.22 | -20.469 | 0.122799 |
| Pose390 | 0.02 | -13.132 | 1.10694 |
| Pose437 | 0.02 | -12.425 | 9.66585 |

The similar method has been applied on mouse FANCD2 to get the monoubiquitinated structure and discussed in next section.

#### 3.3. Optimized structure of human monoubiquitinated FANCD2 and mouse monoubiquitinated FANCD2

To establish the optimized structure molecular dynamic simulation has been performed with the above predicted structure. First step is to do the solvation and to do so explicit periodic boundary (orthorhombic cell shape) model has been opted and it has required ~43,000 water molecules to dissolve the giant structure. Then energy minimization have been done once by steepest decent method and once by conjugate gradient method followed by the heating at 300K for 120ps and equilibration for 200ps. After equilibration the system has been put for dynamic production run for 20ns. The energy of the system and the RMSD have become constant during the production run. Potential energy, backbone root mean square deviation (RMSD), backbone root mean square fluctuation (RMSF), and radius of gyration (Rg) has been analyzed throughout the trajectory to investigate the dynamic performance and stability of the simulated protein. Potential energy vs time has been plotted (**Fig. 7a**) and has been observed that average value of potential energy remains almost constant in the conformers generated between 10 ns - 20ns in MD simulation run. The backbone RMSD profile of optimized monoubiquitinated FANCD2 has been compared with the initially minimized structure. Backbone RMSD have shown fluctuation till 12ns, then it remains almost constant (**Fig. 7b).** Subsequent analysis of root mean square fluctuation (RMSF) of human monoubiquitinated FANCD2 has been illustrated in Fig. 6c, no significant fluctuation (>1 Å) has been observed. The radius of gyration (63.15±0.05Å) (**Fig. S7**) remains almost constant throughout the molecular dynamic simulation run. There has been very small change in radius of gyration for last 10 ns in simulation run. This indicates the structural compactness of the monoubiquitinated FANCD2 complex without structural unfolding. The optimized structure (**Fig. 8**) has been evaluated by Ramachandran Plot, ProQ and Verify3D, and all the data indicates that the overall quality of the predicted structure is compatible with the protein structure of similar size. Solvent accessible surface area has also been evaluated in **Fig. 8b**. The overall trend of potential energy profile, backbone RMSD, RMSF, and radius of gyration for bound and unbound systems have indicated that all systems has been well equilibrated and stable during the simulation run.

**Fig. 7:**
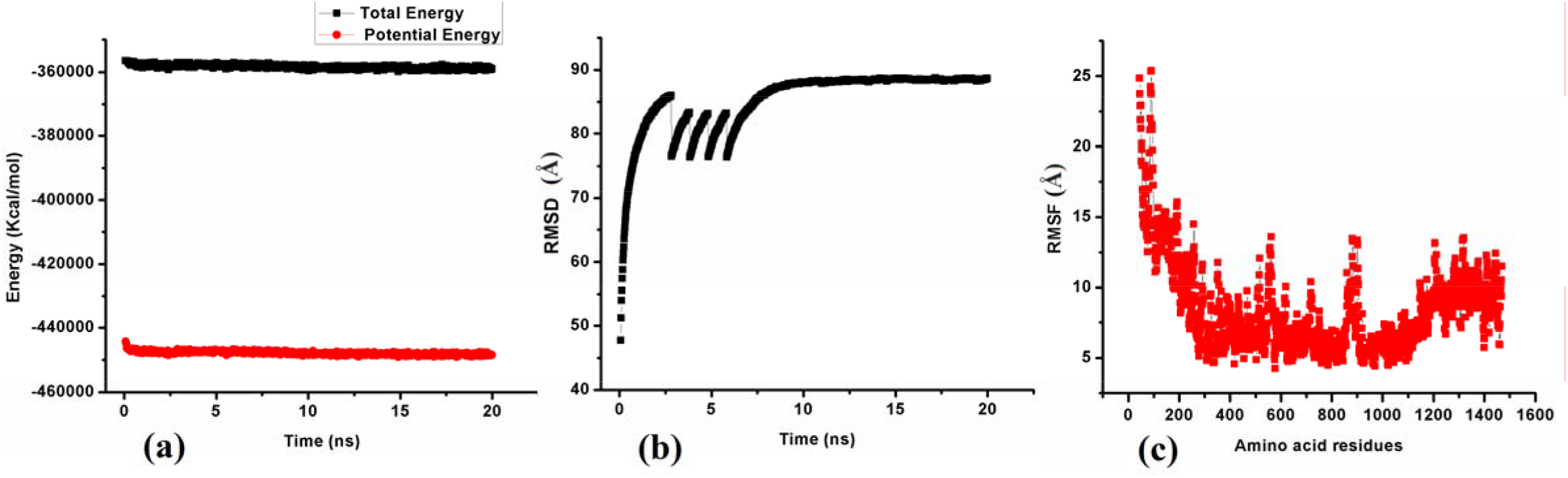
Graphs associated with the molecular dynamic simulation of human monoubiquitinated FANCD2; (a) Energy (Kcal/mol) vs time (ns), (b) RMSD(Å) vs tine (ns) and (c) RMSF vs amino acids residues.

**Fig. 8:**
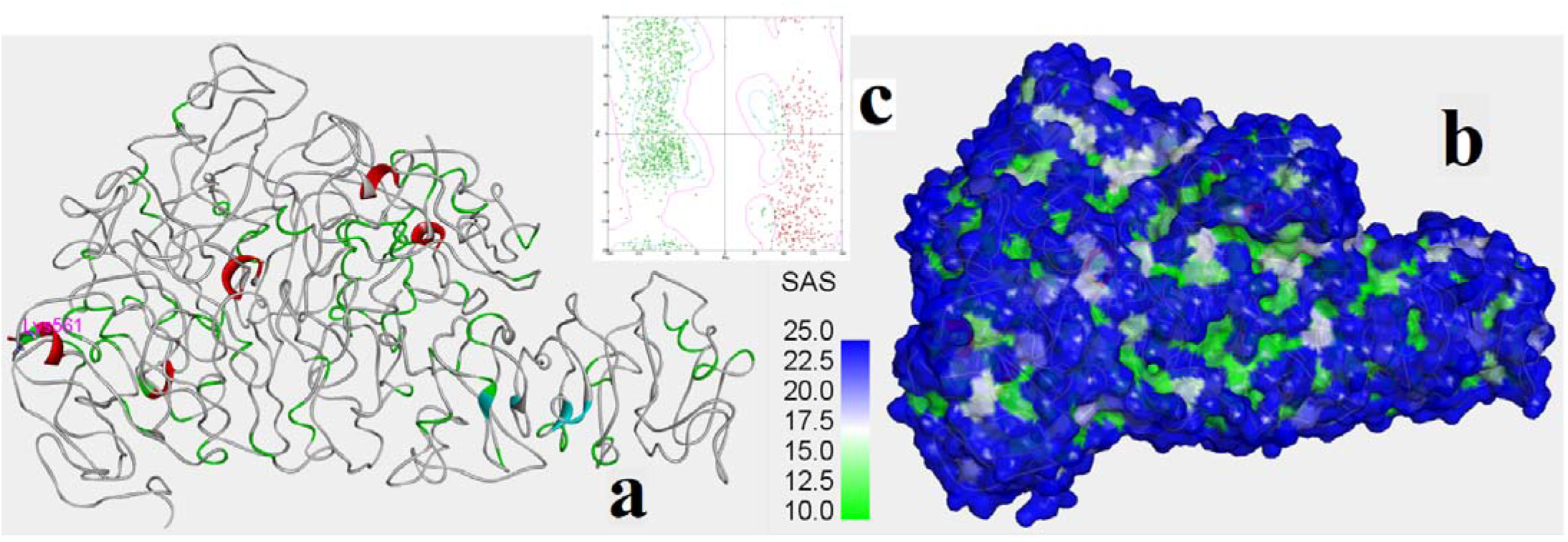
Optimized structure of monoubiquitinated FANCD2 (a), its surface picture (b) and Ramachandran plot (c)

The same procedure has been applied on mouse FANCD2.i.e. protein-protein docking (at Lys559 position) followed by the selection of good poses, then formation of iso-peptide bond in appropriately docked pose and finally molecular dynamic simulation to get the optimized structure of mouse FANCD2. MD simulation has been run till the energy profile, backbone RMSD, RMSF and radius of gyration has reached the constant value. Structure of monoubiquitinated FANCD2 (mouse) before optimization is given in Supplementary file Fig.S8. The optimized structure, MD simulation data and Ramachandran plot are given in **Fig.9, Fig.10, Fig.S9**.

**Fig. 9:**
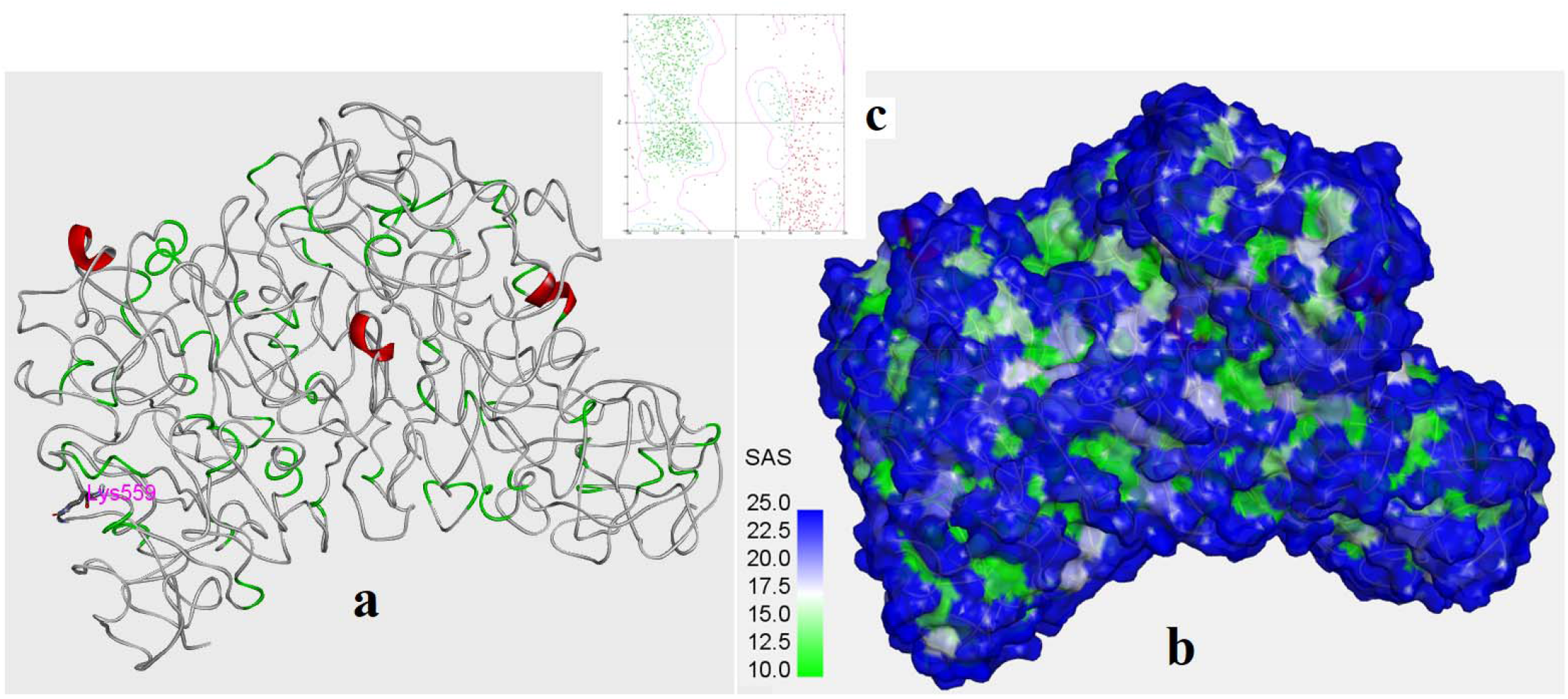
Average structure of monoubiquitinated FANCD2 of mouse (a); and its surface (b); Ramachandran plot (c).

**Fig. 10:**
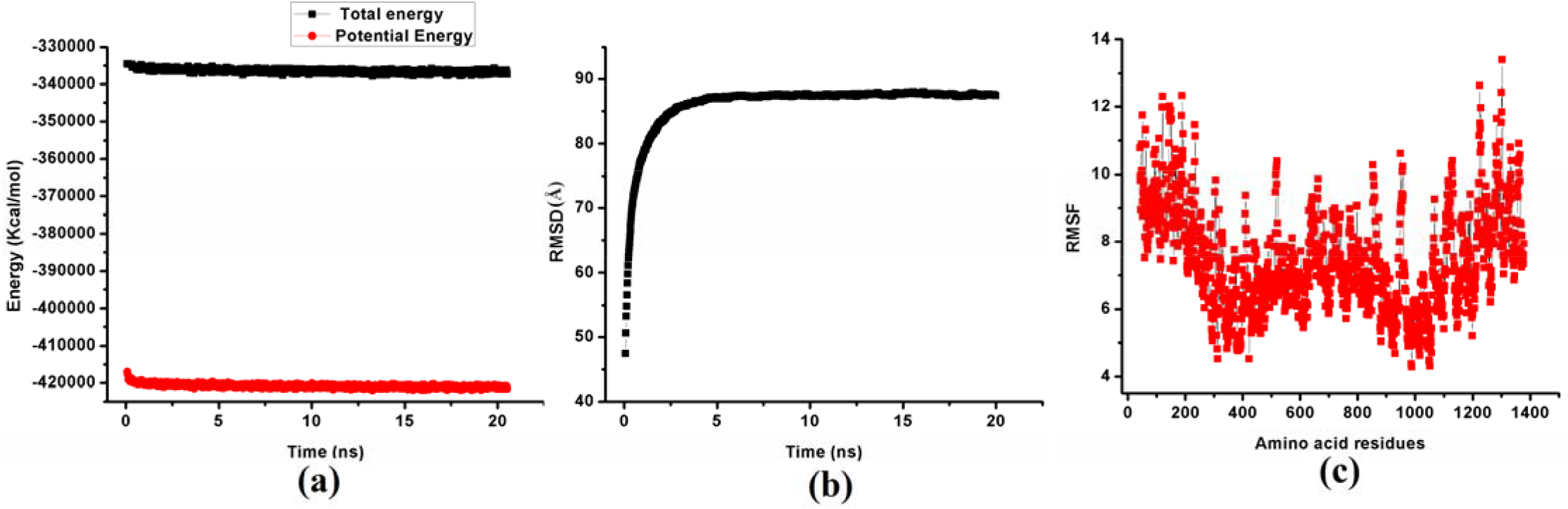
Graphs associated with the molecular dynamic simulation of molecular dynamic simulation of mouse monoubiquitinated FANCD2; (a) energy vs simulation time, (b) RMSD (back bone root mean square deviation) vs time and (c) RMSF (root mean square fluctuation of backbone) averaged over each amino acids.

#### 3.4. Comparison of structure of FANCD2 and monobiquitinated FANCD2

Structure of the FANCD2 and monoubiquitinated FANCD2 is very much different in structure after optimization. Almost similar pattern has been observed in human and mouse system. The FANCD2 remains mostly in solenoid form [**Fig. 2** and crystal structure 3S4W (**Fig. 10a**)]. After optimization of human FANCD2 there are around 50 helices (the majority of the residues stay in helices) while few remain as loop and only 10-12 residues remain as β sheet. The residue of our main interest (Lys561for human and Lys 559 in mouse) remains in a helix. Structure and solvent accessible surface area of the FANCD2 and monoubiquitinated FANCD2 protein (human and mouse) is compared in **Fig.S10 and Fig.11**. In the FANCD2 (human and mouse), the residues mostly remains as helix (**Fig.S10a, Fig.11a**) but after monoubiquitination few residues come out from the helix and form loops and turns, as a result overall helices become less stable (**Fig.S10b and Fig.11b**) and the whole protein turns out to be less compact. Rg value obtained after the MD simulation of FANCD2 and monoubiquitinated FANCD2 supports that prediction. For human FANCD2 the Rg value is 60.7(± 0.5) Å but in case of monoubiquitinated FANCD2 the R_g_ value is 63.15(±0.05) Å (**Fig.S7**). In case of mouse monoubiquitinated FANCD2 the R_g_ value is 62.7(±0.05) Å (**Fig.S9)**. In **Fig.11** apparently it looks that all the helices in **11a** are missing in **11b** but when the Ramachandran plot of FANCD2 (**Fig.S11a**) and monoubiquitinated FANCD2 (**Fig.S11c**) are checked, it is observed that though majority of the residues belong to helical region but few residues that are in helical province in FANCD2 have moved out of the helical area after monoubiquitination, it may be effect of mono ubiquitination (**Fig.11**). Similar effect has been observed when Ramachandran plot of mouse FANCD2 and monoubiquitinated FANCD2 is compared (**Fig.S11**). If we compare Fig.2 and Fig. 5 it is observed that a considerable number of residues in Fig. 2 have been converted to turn and loop in Fig.5. Groban et. al, 2006 reports that conformational change in protein helices and loops are induced by post translational modification. Levy et. al., 2001 also have reported that helix to loop transformation is possible in molecular dynamic simulation in explicit solvent model. So it may be foretold that after mono ubiquitination a structural change is taking place and this structural difference may be responsible for recognising damaged DNA. Our theoretical prediction says that monoubiquitination (at K561 in case of human and K559 in case of mouse) leads to considerable amount of structural change in FANCD2. We can see that FANCD2 in cryo EM structure (6TNG, 6VAD; Alcon et.al, 2020; Wang et.al., 2020) is substantially different from the monoubiquitinated one (6TNF, 6VAF). We also mention that since we are reporting monoubiquitinated FANCD2 (without being docked with DNA) after molecular dynamic simulation (at 300k) and the reported monoubiquitinted structures are already bound to DNA, our computed structure can’t be directly compared with the cryo EM structure. Fig.Sx, Fig.Sx depicts that a considerable amount of structural change has been taken place on monoubiquitination.

**Fig. 11:**
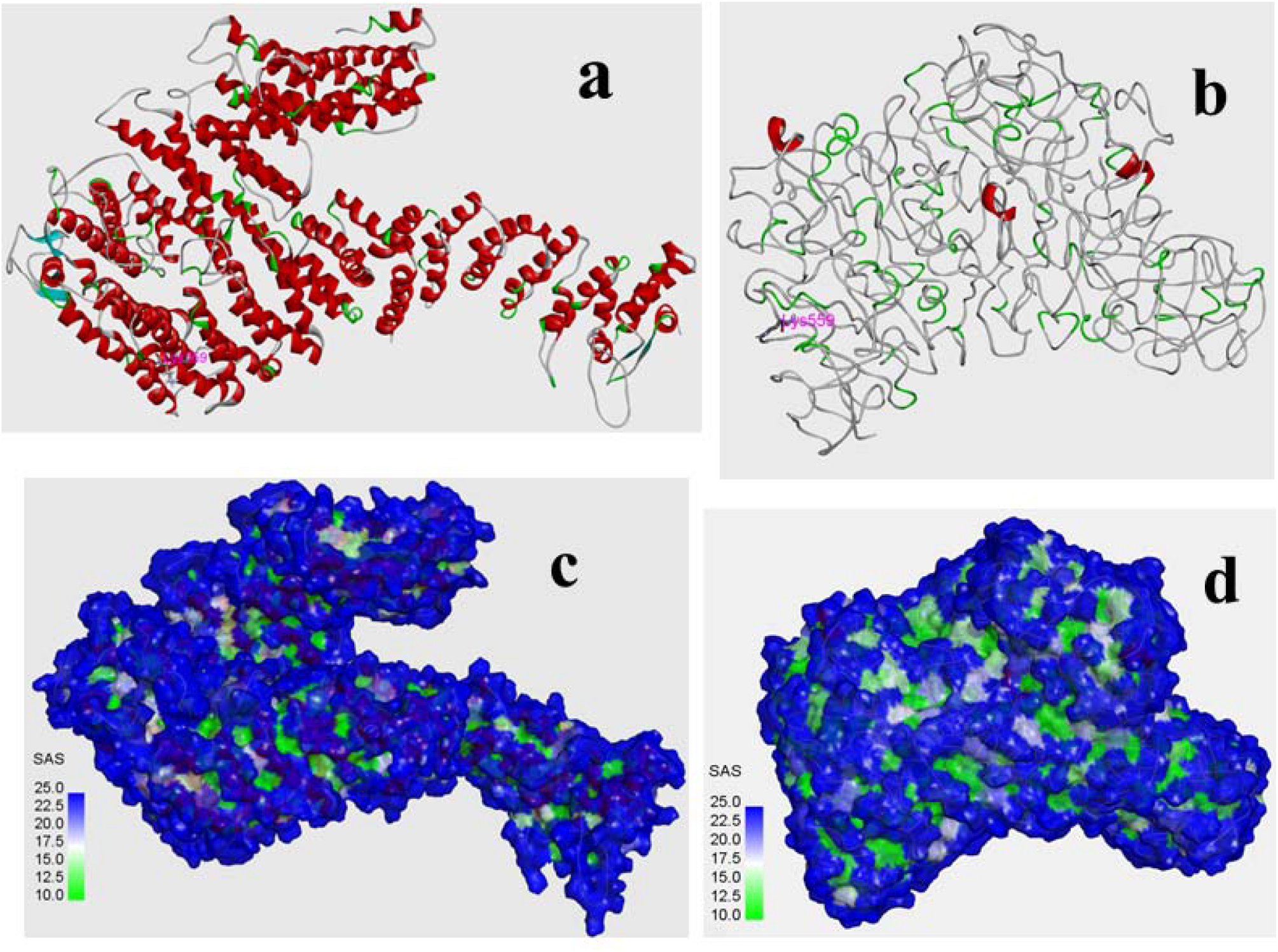
structure of mouse FANCD2 (a), its surface picture (c) and optimized structure of monoubiquitinated FANCD2 of mouse system (b) with its surface picture (d).

**Fig. 12:**
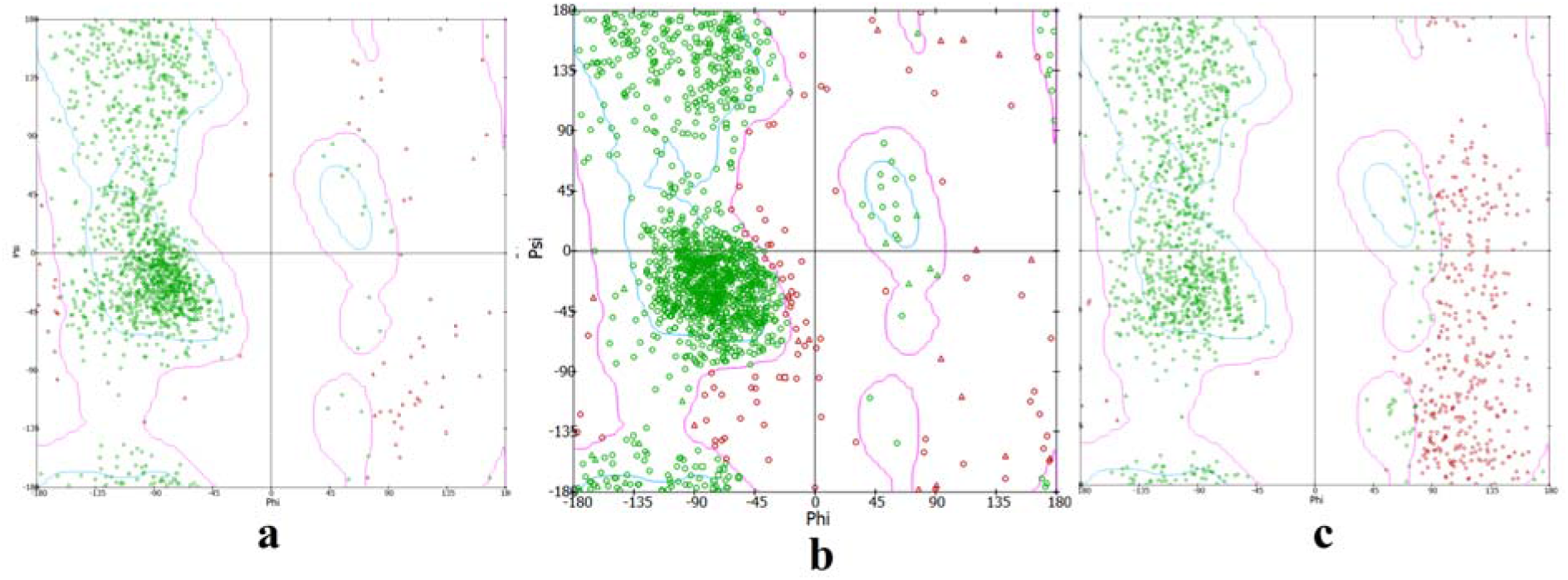
Ramachandran plot of human FANCD2 (a), mono ubiquitinated FANCD2 before simulation (b) and monoubiquitinated FANCD2 after simulation(c).

#### 3.5. Docking of ubiquitinated FANCD2 and DNA

Docking of human FANCD2 and monoubiquitinated FANCD2 with DNA (PDB id 5i50, 6vaa, 6vae, 3aaf) has been done in PatchDock server. Algorithm of PatchDock is based on shape complementary principles. As the FANCD2 and monoubiquitinated FANCD2 has difference in structure and shape, they bind differently with DNA (**Fig.13, S13, S14**). From the docking studies it has been observed that DNA (PDB id 5i50) binds to Arg401, Arg404, Asn 485, Lys828, Asp 849, Asp947, Thr 955, Glu 956, Ser 973, Gln 874, Ile 984, Gln 1053, Tyr 1055, Ser 1059, Gln 1067, His 1356 in case of human FANCD2 **(Fig. 14).** There are salt bridge and H-bond type interactions between the FANCD2 and DNA. Except Asp947 other residues interact with phosphate backbone, Asp947 interacts with O-atom of cytosine. Details of the interaction is given in **Table 2.** In case of monoubiquitinated FANCD2 (human) the double stranded DNA (PDB id 5i50) binds in the region where Lys 190, Lys 261, Lys 248, Lys 1296, Arg 1299, Lys 1361, Asn 257, Ala 291, Lys 358, Cys 432, Glu1303, Leu1360 amino acid residues interact **(Fig 15, Table 3).** All the amino acids except Cys432 interact with the phosphate backbone but Cys432 binds with adenosine. FANCD2 undergoes a structural change upon ubiquitination so FANCD2 and monoubiquitinated FANCD2 binds differently with same DNA sequence. We have repeated the docking studies with other DNA sequences (DNA structure available in the cryo EM structures with PDB id 6VAA, 6VAE and crystal structure with PDB id 3AAF) and have obtained similar result. When the DNA sequence available in the cryo EM structure (PDB id 6VAE) binds to human FANCD2 it interacts with Lys358, Lys397, Arg401, Ala487, Glu488, His853, Gly909, Ala923, The952, Arg987, Gln1067 by the formation of H-bond and salt bridge (**Fig.16a, Table S1**); but when binds to monoubiquitinated FANCD2 it interacts with Lys190, Lys261, Lys397, Ser433, Ser434, Val512, Lys1248, Lys1296, Arg1299, Lys1361 (**Fig.16b, Table S2**). Here Lys387 forms H-bond with the nucleotide (thymine) and His853 forms H-bond with pyranose ring of backbone while interacting with human FANCD2, all other amino acids interact through phosphate backbone of DNA. Similarly the DNA sequence available in PDB id 6VAA binds to Arg401, Gln823, Lys828, Arg986, Arg1299, Glu827, His853, Asn920, Gln974, Lys1051, Gln1053, Gln1067, His1356 when it fixes with human FANCD2 structure but it networks with Arg253, Thr290, Arg355, Asp400, Cys432, Lys1296, Arg1299 while it is attached to monoubiquitinated FANCD2 (**Fig.S15, Table S3,S4**). In case of the DNA sequence present in PDB id 3AAF the residues responsible for interaction with human FANCD2 are Gln823, Lys828, Ile984, Arg986, Gln1053, His1056, Ser1059, Gln1067 and the residues liable for fixing with monoubiquitinated FANCD2 are Asn257, Lys261, Thr290, Arg355, Lys397, Ser434, Lys1248, Lys1296, Arg1299, Glu1303, His1356, Leu1360 (**Fig.S16, TableS5, S6**).

**Fig. 13:**
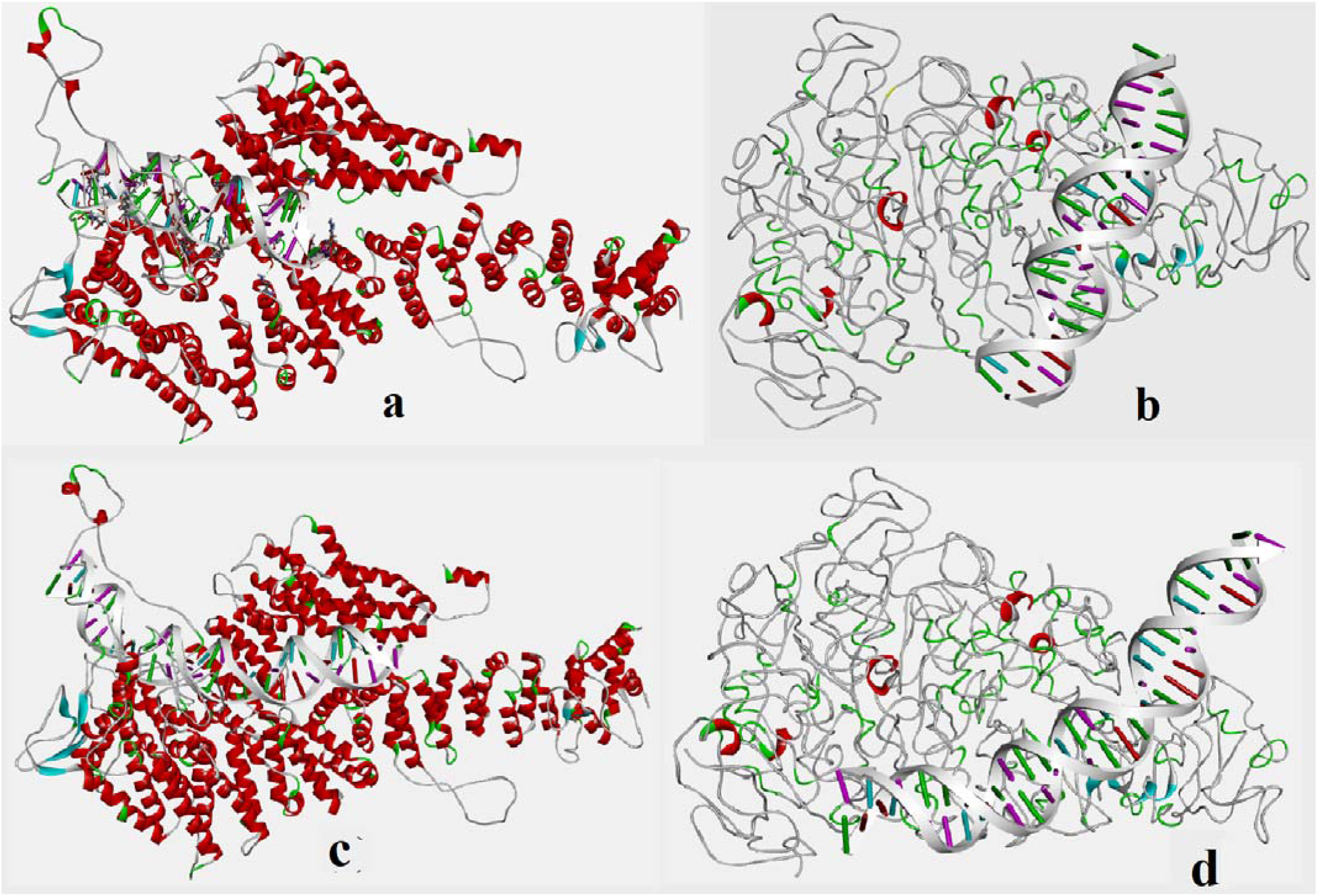
DNA (PDB id 5i50) docked with Human FANCD2(a) and monoubiquitinated FANCD2 (b); DNA from 6VAE is docked with human (c) FANCD2, (d) monoubiquitinated FANCD2.

**Fig. 14:**
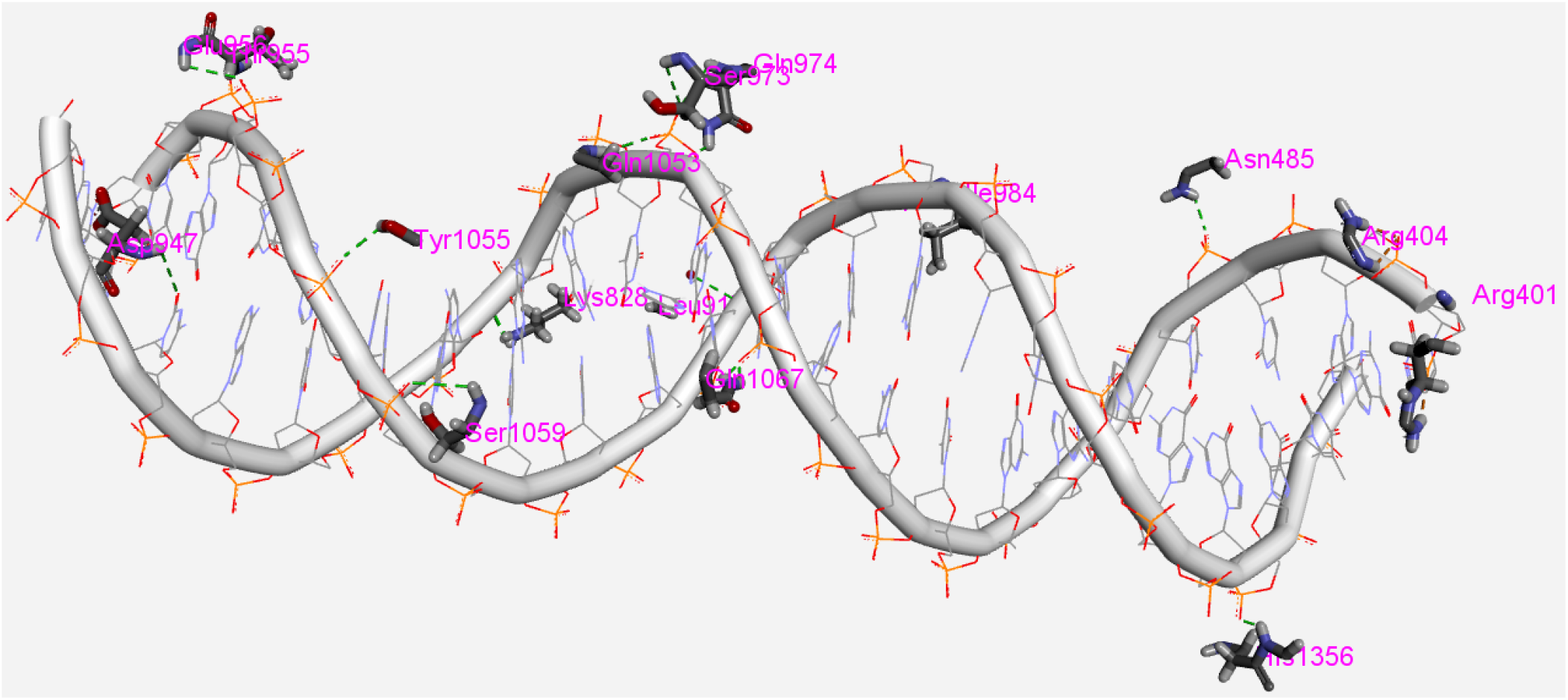
Close view of the inte_r_action when DNA (5i50) is docked with human FANCD2.

**Fig 15:**
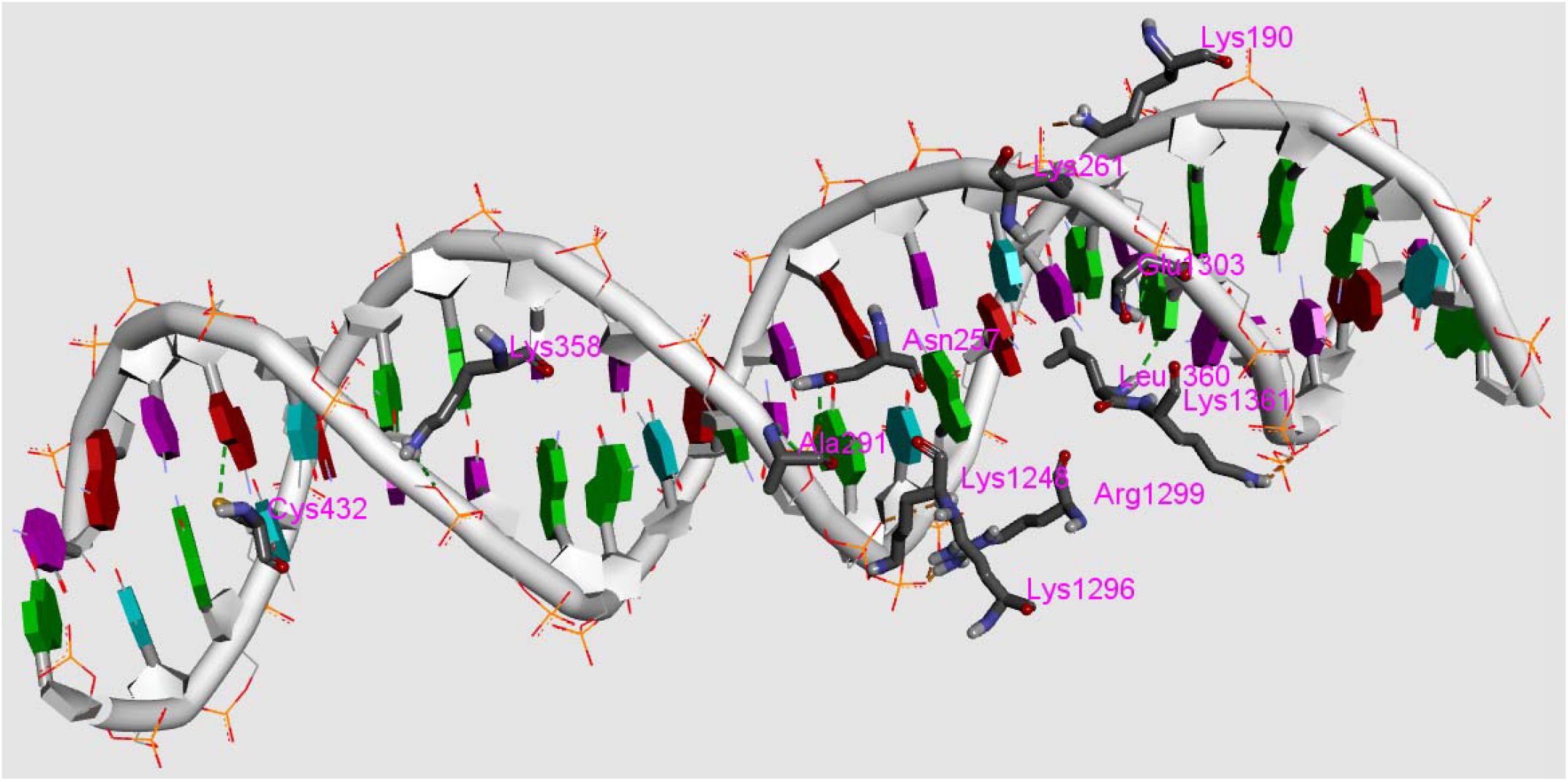
Close view of the interaction of DNA (PDB id 5i50) and monoubiquitinated FANCD2 (human) in best docked pose.

**Fig. 16:**
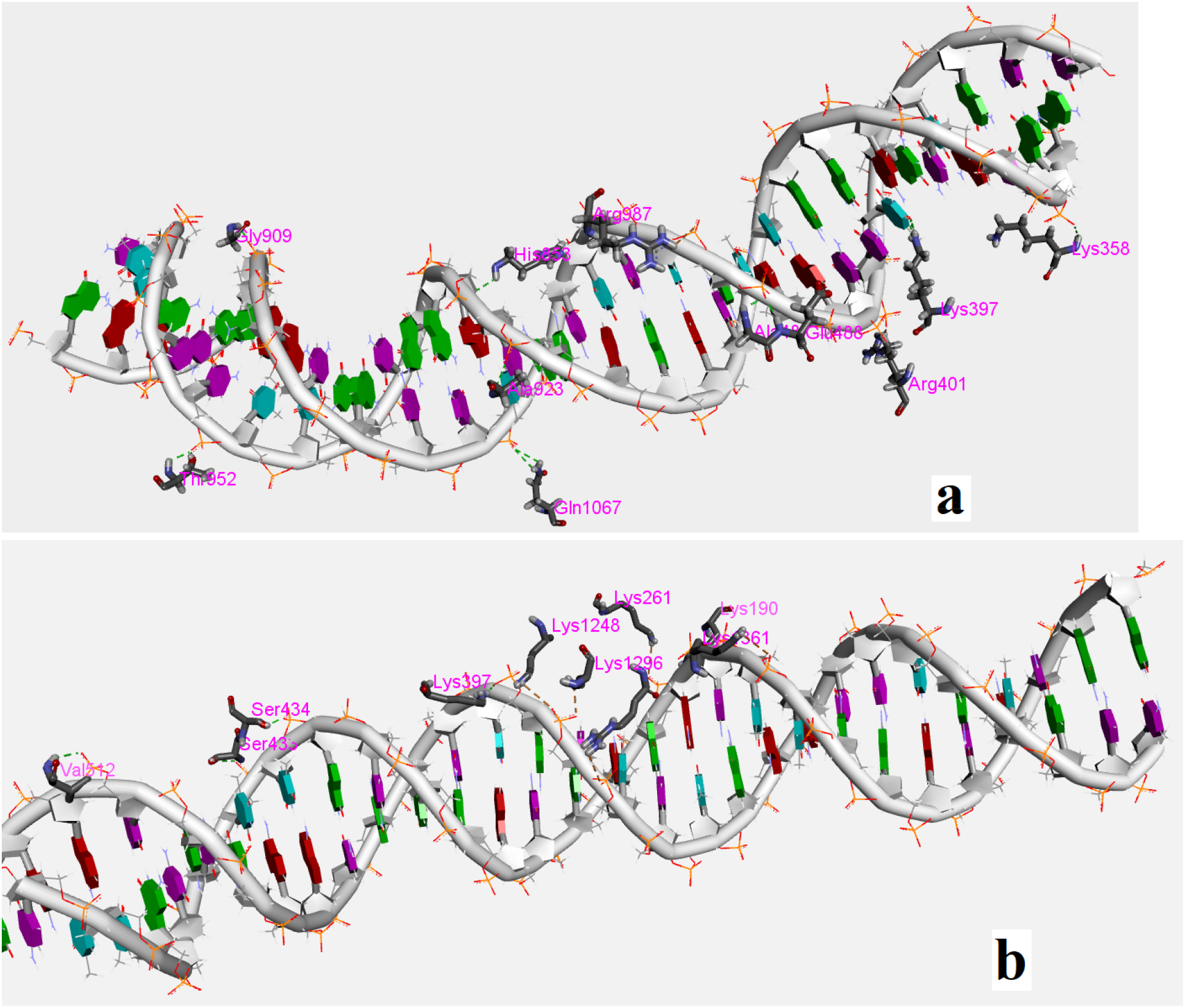
Close view of the interaction of DNA (PDB id 6VAE) with human FANCD2 **(a)** and monoubiquitinated FANCD2 (human) **(b)** in best docked pose.

**Table 2:**
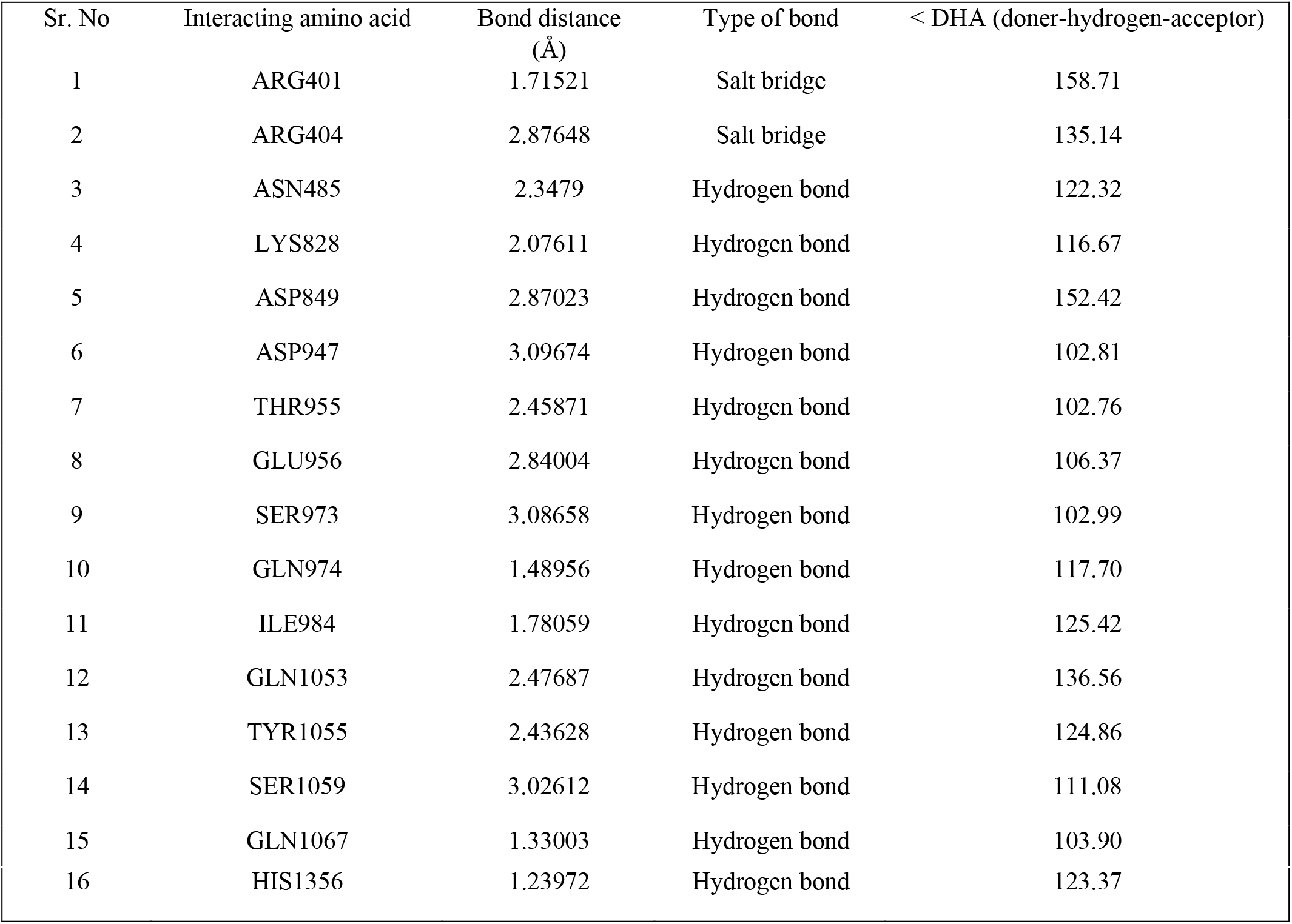
Details of the interaction when DNA (PDB id 5I50) is docked with humanFANCD2.

| Sr. No | Interacting amino acid | Bond distance (Å) | Type of bond | < DHA (doner-hydrogen-acceptor) |
| --- | --- | --- | --- | --- |
| 1 | ARG401 | 1.71521 | Salt bridge | 158.71 |
| 2 | ARG404 | 2.87648 | Salt bridge | 135.14 |
| 3 | ASN485 | 2.3479 | Hydrogen bond | 122.32 |
| 4 | LYS828 | 2.07611 | Hydrogen bond | 116.67 |
| 5 | ASP849 | 2.87023 | Hydrogen bond | 152.42 |
| 6 | ASP947 | 3.09674 | Hydrogen bond | 102.81 |
| 7 | THR955 | 2.45871 | Hydrogen bond | 102.76 |
| 8 | GLU956 | 2.84004 | Hydrogen bond | 106.37 |
| 9 | SER973 | 3.08658 | Hydrogen bond | 102.99 |
| 10 | GLN974 | 1.48956 | Hydrogen bond | 117.70 |
| 11 | ILE984 | 1.78059 | Hydrogen bond | 125.42 |
| 12 | GLN1053 | 2.47687 | Hydrogen bond | 136.56 |
| 13 | TYR1055 | 2.43628 | Hydrogen bond | 124.86 |
| 14 | SER1059 | 3.02612 | Hydrogen bond | 111.08 |
| 15 | GLN1067 | 1.33003 | Hydrogen bond | 103.90 |
| 16 | HIS1356 | 1.23972 | Hydrogen bond | 123.37 |

**Table 3:** Details of the interaction when DNA (PDB id 5I50) is docked with human monoubiquitinated FANCD2.

| Sr. no | Interacting amino acid | Bond distance (Å) | Type of bond | < DHA (doner-hydrogen-acceptor) |
| --- | --- | --- | --- | --- |
| 1 | LYS190 | 1.914 | Salt Bridge | 120.57 |
| 2 | LYS190 | 1.998 | Salt Bridge | 135.14 |
| 3 | LYS261 | 3.217 | Salt Bridge | 154.07 |
| 4 | LYS1248 | 1.46 3 | Salt Bridge | 120.77 |
| 5 | LYS1296 | 2.771 | Salt Bridge | 127.09 |
| 6 | ARG1299 | 0.964 | Salt Bridge | 138.95 |
| 7 | ARG1299 | 1. 346 | Salt Bridge | 136.04 |
| 8 | LYS1361 | 2.927 | Salt Bridge | 127.90 |
| 9 | ASN257 | 2.101 | Hydrogen Bond | 115.89 |
| 10 | ALA291 | 2.218 | Hydrogen Bond | 117.40 |
| 11 | LYS358 | 2. 342 | Hydrogen Bond | 136.60 |
| 12 | CYS432 | 3.692 | Hydrogen Bond | 112.80 |
| 13 | GLU1303 | 2.202 | Hydrogen Bond | 94.26 |
| 14 | LEU1360 | 2. 338 | Hydrogen Bond | 159.1 |

## 4. Conclusion

Here we have modelled the structure of monoubiquitinated FANCD2 and found that structure of FANCD2 is significantly different from the monoubiquitinated FANCD2. FANCD2 is a very big protein and doing molecular dynamic simulation of such a big protein was very much challenging. We have overcome the challenge and have managed to run the simulation until it becomes energetically stable and has reached the constant RMSD value. From the theoretical result we have predicted that a structural change is taking place after monoubiquitination. We have also predicted a method to do monoubiquitination theoretically by protein-protein docking followed by molecular dynamic simulation. We are extending our work with FANCD2-FANCI heterodimer and trying to unfold the mystery of Fanconi anemia DNA damage repairing pathway.

## 5. Acknowledgement

I acknowledge SERB for the financial support (PDF/2016/002799). I also acknowledge CSIR. I am thankful to NIT Durgapur for providing me the work space. I must acknowledge Dr. Ankur Chaudhuri, research associate of State University, Barasat and Anand Krishnamurthy of Dassult System for their advice for running protein-protein docking and molecular dynamic simulation of such a big protein.

